# Rapid Color Categorization in the Brain Revealed by Frequency-tagging-based EEG

**DOI:** 10.1101/2022.07.28.501827

**Authors:** Mengdan Sun, Xiaoqing Gao

## Abstract

The origin of color categories has been debated extensively. Historically, linguistic relativists claim that color categories are shaped by the language we speak and that color terms subsequently affect our perception of color, while universalists postulate that color categories are independent of language and formed based on perceptual mechanisms. A recent hypothesis suggests that the original fine-grained color space in the visual cortex may be transformed into categorical encoding due to top-down modulation. To test the nature of color categorization, our study adopted a sensitive frequency-tagging-based EEG paradigm where the color stimuli were presented sequentially at a fast speed of 10 Hz (SOA: 100 ms) to probe fast, implicit processing of color categories. This SOA was supposed to disrupt top-down feedbacks in visual processing. We showed that EEG responses to cross-category oddball colors at the frequency where the oddball stimuli were presented was significantly larger than the responses to within-category oddball colors. This finding suggested that the brain encodes color categories automatically when top-down feedbacks from frontoparietal areas are blocked. Our study supports the view that the categorical processing of color emerges at the early perceptual stage.

## 1. Introduction

The spectrum of color is continuous and fine-grained, yet it is described by only several discrete categories (e.g., green, blue) in our language. The origin of color categories has been debated extensively (for review, see Regier & Kay, 2009; Siuda-Krzywicka et al., 2019a; Witzel, 2019). Historically, linguistic relativists claim that color categories are shaped by the language we speak and that color terms subsequently affect our perception of color, while universalists postulate that color categories are independent of language and formed based on perceptual mechanisms. More recent evidence contradicts a clear-cut dichotomy between relativism and universalism. It has been agreed that color categories are shaped by both universal constraints and language (Abbott et al. 2016; Kay & Regier 2006; Kuehni 2007; Regier & Kay 2009; Roberson & Hanely, 2009; 2010). A recent hypothesis about the nature of color categories suggests that the original fine-grained color space in the visual cortex may be transformed into categorical encoding due to top-down modulation (e.g., Brouwer & Heeger, 2013; Lupyan et al., 2020; Kwok et al., 2011; Siok et al., 2009; Sun et al., 2021; Witzel, 2019). The top-down influence could derive from high-level cognitive processes such as language or (and) attention. This hypothesis features an online warping of the perceptual color space rather than a permanent change. But the top-down hypothesis still needs to be systematically tested.

Electroencephalography (EEG) has been repeatedly adopted to understand the nature of color categories (Athanasopoulos et al., 2010; Clifford et al., 2010; 2012; Fonteneau & Davidoff, 2007; Forder et al., 2017; He et al., 2014; Holmes et al., 2009; Mo et al., 2011; Thierry et al., 2009; Zhong et al., 2015), as the event related potentials (ERP) modulated by color categories provide rich information about the processing stages on which the category effects emerge. For instance, both P1 (Fonteneau & Davidoff, 2007; Holmes et al., 2009; Maier & Rahman, 2018) and Visual Mismatch Negativity (vMMN) (Athanasopoulos et al., 2010; Clifford et al., 2010; Mo et al., 2011; Thierry et al., 2009; Zhong et al., 2015) have been found to be involved in color categorization, suggesting that early, pre-attentive color perception is shaped by color categories. It is noteworthy that some findings are subjective to insufficient stimulus control (for review, see Siuda-Krzywicka et al., 2019a; Witzel, 2019), and therefore, the observed category effect may not be genuine. On the other hand, some studies suggested that color categories can only modulate late stages of visual processing, indexed by P2 (Forder, 2015), P3 (Clifford et al., 2012; He et al., 2014), and N2 (Forder, 2015; He et al., 2014; Liu et al., 2009). There are also studies showing the presence of both early and late components in category effects (Fonteneau & Davidoff, 2007; Holmes et al., 2009; Maier & Rahman, 2018).

Despite the possibility of insufficient stimulus control (Siuda-Krzywicka et al. 2019a; Witzel & Gegenfurtner, 2015), the inconsistency in previous EEG findings may result from the variability in the experimental procedures. First, the previous studies differed in the task requirements for participants. In some studies, the participants were explicitly required to make color judgments (e.g., count the number of deviant color stimuli among a stimulus sequence in Clifford et al., 2012 and He et al., 2014). Other studies used passive viewing tasks to probe implicit and automatic color processing (Clifford et al., 2010; Forder et al., 2017; Thierry et al., 2009), in which the participants’ attention was directed away from the color stimuli. Explicit color judgment may introduce attention, verbal coding, or decisional processing into the EEG responses, consequently affecting the stages in which category effects may occur. In addition, the stimulus onset asynchronies (SOA) were varied in the previous studies, ranging from 94 ms (Maier & Rahman, 2018) to about 2,000 ms (Clifford et al., 2012). The SOAs place constraints on the types of neural processes involved in visual recognition (Lamme & Roelfsema, 2000; Potter et al., 2014). For example, top-down feedback processing from the frontoparietal cortex is disrupted during short SOAs. Thus, the varying SOAs might also cause inconsistency in previous findings.

To probe fast, implicit processing of color categories, the current study adopted a frequency-tagging-based EEG paradigm to examine the nature of color categorization. Our paradigm resembles the rapid serial visual presentation (RSVP) paradigm, which is widely used to study the dynamic temporal properties of human perception and cognition (Fabre-Thorpe et al., 2001; Potter & Levy, 1969; Potter et al., 2014). The processing time of visual stimuli in our study was strictly controlled because the stimuli were both pre- and post-masked. In our paradigm, color stimuli were presented at a fast speed of 10 Hz, corresponding to a short SOA of 100 ms. A review by Wyatte et al. (2014) suggested that the earliest top-down feedbacks are engaged at about 130 ms after stimulus onset. Therefore, an SOA of 100 ms combined with backward masking can disrupt the involvement of feedback loops from frontoparietal areas (Lamme & Roelfsema, 2000; Potter et al., 2014; Wyatte et al., 2014). If top-down modulation from high-level cognitive processes is necessary for color categorization in the human brain, then we would expect the absence of category effects under the SOA of 100 ms. If categorical encoding of color originates from the visual system, then we would observe category effects under such SOA. It is noteworthy that there are local recurrent feedbacks within the visual hierarchy (from higher-level visual areas to lower-level areas). These local feedbacks emerge earlier than top-down feedbacks from frontoparietal areas, which is not our research interest here.

In addition, we implemented frequency tagging (Adrian & Matthews, 1934; Regan, 1989; Norcia et al., 2015) in the RSVP paradigm. Among the 10-Hz RSVP stream (i.e., base frequency), oddball color stimuli were periodically presented at 2 Hz (i.e., oddball frequency), and multiple baseline (or standard) color stimuli were inserted in between every two oddball stimuli. Compared with the traditional ERP paradigm, the frequency-tagging EEG paradigm has a greater signal-to-noise ratio (SNR) and sensitivity (Adrian & Matthews, 1934; Regan, 1989; Norcia et al., 2015; Rossion, 2014), which helps us to probe subtle neural response to very brief color stimuli. To exclude neural processes caused by explicit color judgment while keeping the participants attending to the color stimuli, we asked participants to monitor changes in fixation cross overlayed on the color stimuli, without responding to the color stimuli. The oddball and the baseline colors were either from the same category (within-category) or two different categories (cross-category), with perceptual differences equalized. The EEG response at the oddball frequency would reflect a differential response between the base color and the oddball color, providing a direct measure of how the brain discriminates the oddballs from the base stimuli. If the brain categorizes color in the current experimental settings, then the response at the oddball frequency in the cross-category condition would be greater than the response in the within-category condition. Otherwise, there would be no significant difference between the two conditions.

## 2. Methods

### 2.1 Participants

Seventeen participants (8 males, mean age = 22.53, SD = 2.72) with normal or corrected to normal vision were recruited from Zhejiang University. They all signed informed consent forms and received monetary compensation for their participation. None reported any history of psychiatric or neurological disorders. This study was approved by the Research Ethics Board of the Center for Psychological Sciences at Zhejiang University (certificate NO. 2020-008).

### 2.2 Stimuli

The color stimuli were checkerboards contrasting a color and a neutral background (Fig.1A). The checkerboards consisted of a 10 × 10 grid of square checks and subtended approximately 10.6° × 10.6° of visual angle. For each checkboard, a random half of the square checks was specified by one of the six colors. The six colors used in the checkerboards were chosen from CIELAB space (L*=70, a radius of 29) which only varied in hue (Fig.1B). The hue degrees of these six stimuli are 140° (G1), 170° (G2), 180° (G3), 200° (B1), 210° (B2), and 240° (B3). The category membership of each color stimulus was determined by a green-blue boundary (192.1°) defined from a preliminary color naming experiment. Note that the participants in the preliminary color naming experiment were not recruited in the subsequent EEG experiment. The luminance of the background gray was equal to that of the colors.

**Fig. 1.**
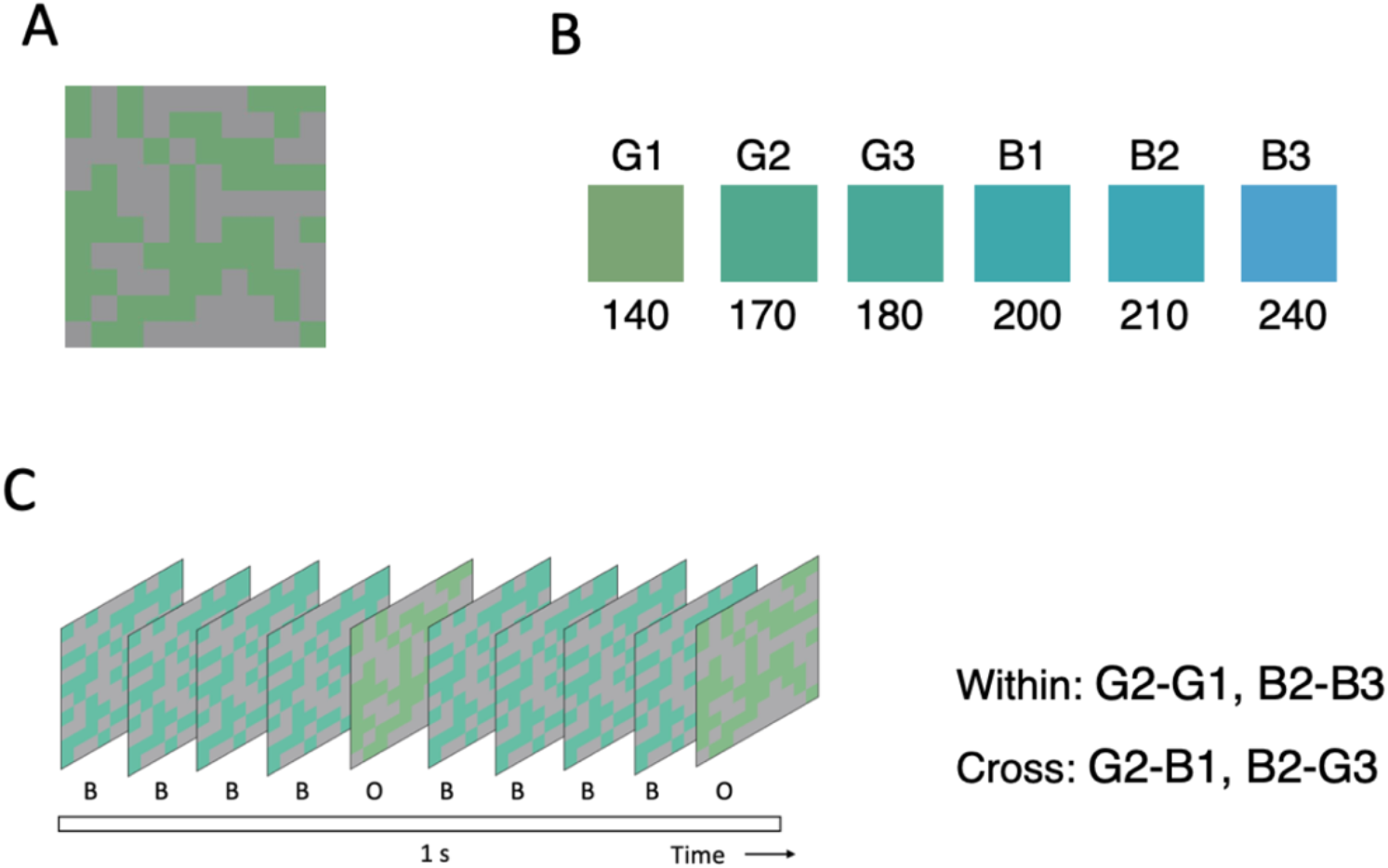
Stimuli and procedure. (A) Example of color stimuli. (B) The six colors used in the checkerboards were chosen from CIELAB space (L*=70, a radius of 29), which only varied in hue. (C) Stimuli were presented at a rate of 10 Hz (1 cycle = 100 ms). In each stimulation sequence, the oddball stimuli (O) were presented at 2 Hz, and four baseline stimuli (B) were presented in between every two oddballs.

The stimuli were projected onto a 25-inch monitor (screen resolution of 1920 × 1080 at a frame rate of 240 Hz) in a dimly lit room. The Monitor was calibrated using a Minolta Konica cl-500A colorimeter (https://sensing.konicaminolta.us). The xyY value of the white point (x= 0.3287, y= 0.3505, Y= 137.8) and the monitor primaries (R: x= 0.6297, y= 0.3355, Y= 30.3; G: x= 0.3207, y= 0.6182, Y= 100.7; B: x= 0.1614, y= 0.0764, Y= 11.7) were measured. The inputoutput value of each channel was also measured to define the gamma curve. This information was used to determine the appropriate RGB values for each color stimulus as suggested by Brainard, Pelli, and Robson (2002).

### 2.3 Procedure

#### EEG experiment

During the stimulation, the color stimuli, appeared at the center of a uniform light grey background extending on approximately 33° × 25° of visual angle. The color stimuli were presented at a fixed rate of 10 Hz (base stimulation frequency: 10 color stimuli per second, SOA: 100 ms). Note that there was no interstimulus interval in our paradigm, the SOA is equivalent to stimulus presentation time. Each stimulation sequence was composed of two colored checkerboards, with one colored checkerboard as the baseline stimulus and the other colored checkerboard as the oddball stimulus. The oddball stimuli were presented at a fixed speed of 2 Hz (i.e., oddball frequency). In addition, each stimulation sequence began with a 2 s fade-in period in which the stimuli gradually increased from 0 to 100% contrast and ended with a 2 s fade-out period in which the stimuli decreased from 100 to 0% contrast. The whole stimulation lasted 124 s (including 2 s fade-in and 2 s fade-out). While monitoring the continuously flashed color stimuli, participants were asked to fixate on a central fixation cross and press the SPACE key whenever the fixation changed from black to white (11 change occurred in each stimulation sequence, at random intervals).

The experiment included two conditions in which the oddball and the baseline color were either from the same category (within-category) or different categories (cross-category) (Fig.1C). The within-category condition included two different types of color pairs: G2-G1 (170°, 140°), and B2-B3 (210°, 240°). The cross-category condition included two different types of color pairs: G2-B1 (170°, 200°) and B2-G3 (210°, 180°). Note that the baseline colors in the two conditions were identical (i.e., G2 and B2). Each condition included four 124-s stimulation sequences (two for each color pair). The sequence order was randomized across conditions.

#### Behavioral color naming

After the EEG experiment, the participants were required to complete a color naming task to determine each participant’s subjective green-blue boundary. On each trial, a color stimulus (Fig.1A) was present at the center of the screen. The stimulus was filled with one of the 11 colors (140° to 240°, step size: 10°). The participants needed to respond which color term (green or blue) most closely described the color by pressing the key *F* or *J.* Each participant completed eight trials for each color, with a total of 88 trials presented in a random order.

### 2.4 EEG recording

EEG was recorded with 64 active electrodes mounted in a cap (Easycap GmbH, Brain Products, Munich, Germany) according to the standard 10-20 system, with FCz as reference and AFz as ground. Eye movements were monitored using one electrode placed at the outer canthi of the right eye. EEG was sampled at 500 Hz. Impedance was reduced below 20 kΩ for each electrode by injecting electrode gel.

### 2.5 EEG analysis

#### Preprocessing

All EEG preprocessing steps were carried out using Letswave 5 (http://nocions.webnode.com/letswave) on MATLAB R2014a (The MathWorks, USA). EEG data was first band-pass filtered at 0.1-100 Hz using Butterworth band-pass filter with fourth-order and zero-phase and then downsampled to 250 Hz to reduce data size and computational load. Data were then segmented into 128-s segments, including 2 s before and after each stimulation sequence (−2-126 s). All 64 EEG channels were then re-referenced to the average of all channels except for the ocular channel.

#### Frequency domain analysis

Preprocessed EEG data was further segmented into an integer number of 2-Hz cycles beginning 2 s after the fade-in (to avoid any contamination by the fade-in and initial transient responses), until 120 s after the fade-in (240 cycles, 30,000 time-bins in total). The resulting epochs were averaged for each condition (within-category and cross-category). A discrete Fourier transformation (FFT) was applied to these averaged epochs, and amplitude spectra were extracted for all channels. It resulted in spectra ranging from 0 to 250 Hz with a frequency resolution of 0.0083 Hz (1/120 s). To determine relevant harmonics for the base and oddball frequency, grand-averaged amplitudes were calculated by averaging individual amplitude spectra within each condition. The grand-averaged amplitudes were further averaged across all channels and transformed into z-scores. The z-scores were calculated by subtracting the mean amplitude of 20 neighboring frequency bins (10 bins on each side while excluding the immediately adjacent bins) from that frequency, and then divided by the standard deviation of the 20 neighboring bins. The number of harmonics selected was constrained by the highest channel-averaged z-score that was significant at least in one condition (z-score > 2.32, *p* < 0.01).

The responses at the frequencies of interest were quantified by applying a baseline correction to the individual amplitude spectra, i.e., subtracting from each selected frequency the mean amplitude from the 20 surrounding frequency bins (10 bins on each side while excluding the immediately adjacent bins). Based on previous studies revealing that higher-level visual areas may encode color categories (Brouwer & Heeger, 2013; Sun et al., 2021), we defined occipitotemporal areas as regions of interest (ROIs). In addition, our study was interested in the top-down influence, we also selected frontal areas as ROIs. Thus, bilateral OT (occipitotemporal) electrodes (left: P3, P5, P7, PO3, PO7; right: P4, P6, P8, PO4, PO8) and frontal electrodes (left: F1, F3, F7, AF3, AF7; right: F2, F4, F8, AF4, AF8) were defined as ROIs to assess the effect of color categories. The responses on the five electrodes from each ROI region were averaged to obtain ROI-level responses.

#### Time domain analysis

Complementary to the frequency analysis, we also analyzed the EEG data in the time domain. The re-referenced data were low-pass filtered using FFT at 30 Hz with a cutoff width of 1 Hz. A multi-notch narrowband FFT filter with 0.1 Hz width was used to selectively remove the dominating base frequency and its first four harmonics (10 to 40 Hz). The filtered data was cropped into smaller epochs of 2 s, containing 20 base stimulation cycles. Each epoch contains four oddball cycles (BBBBOBBBBOBBBBOBBBBO). The epochs were averaged separately for each participant and for each condition, and then baseline-corrected relative to the base stimulus presentation preceding the first oddball. Grand-averaged epoch was obtained by averaging individual waveforms for the display of time-domain data.

To determine which time windows showed significant effects of color discrimination, we also ran cluster-based permutation t-tests on the post-stimulus onset time-points within each occipitotemporal region for each condition (0-800 ms, all other parameters same as above; corrected significance level*p* < 0.05; permutation times: 1,000). This analysis was done on baseline-corrected waveform amplitudes.

## 3. Results

### 3.1 Behavioral color naming

The proportion of “blue” responses in the color-naming task was fitted with a sigmoid function defined as 1/ (1 + exp (– (x – α)/β), where α is the threshold (estimated value at which “blue” would be reported half of the time), that is, the green-blue boundary in our study. The participants’ average behavioral green-blue boundary was 187.5° (SD = 7.5°), confirming that the predefined categorical membership is appropriate at the group level for the color stimuli used in the EEG study.

### 3.2 EEG: frequency domain

#### Base frequency

Both conditions showed robust responses at the base stimulation frequency and multiple harmonics (Fig. 2A). The responses at the fourth harmonic remained significant in the cross-category condition (z = 4.41, p < 0.001).

**Fig. 2.**
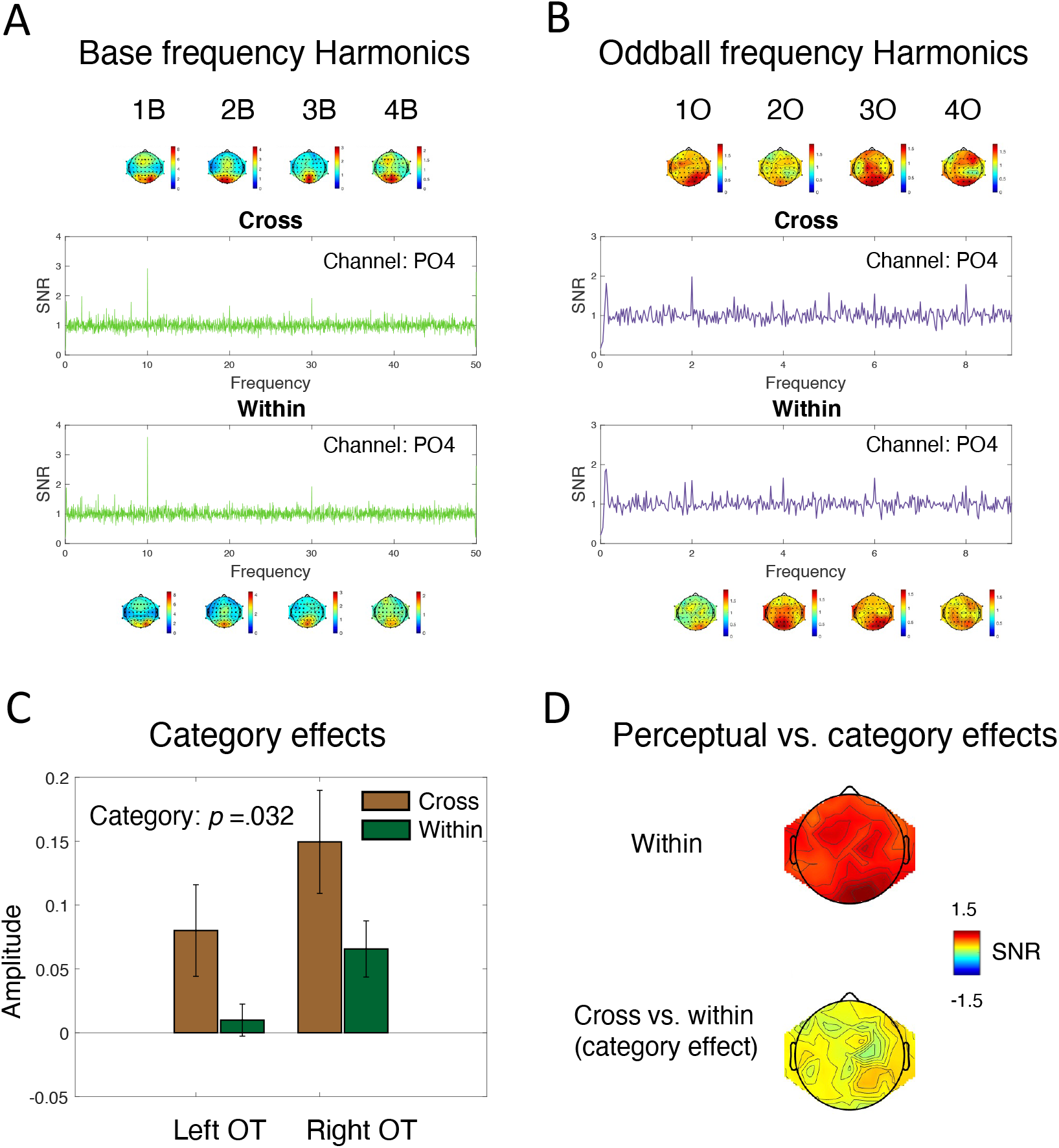
Results of frequency domain analyses. Channel-averaged SNR at the base frequency and its harmonics (1B, 2B, 3B, 4B) (A) and the oddball frequency and its harmonics (1O, 2O, 3O, 4O) (B) in the cross-category and within-category conditions with scalp topographies. The color scale is adjusted to zero and maximum values of each frequency. (C) Signal amplitudes at the oddball frequency in left and right OT regions. Error bars represent 95% CI. (D) Topographic map showing responses to within color stimuli and category-selective responses (cross minus within) at the oddball frequency.

#### Oddball frequency

In addition, we observed significant responses at the oddball frequency (2 Hz) and its five harmonics (4 Hz, 6 Hz, 8 Hz, 12 Hz, and 14 Hz) (Fig. 2B), suggesting neural bias to perceptual color differences (i.e., hue). Overall, the responses at both base frequencies and oddball frequencies were greater in the cross-category condition than in the within-category condition. The responses were peaking around occipitotemporal regions (e.g., PO3, PO4).

To investigate whether the neural activities were sensitive to categorical differences, we averaged the baseline-corrected amplitudes on five electrodes within each ROI (left OT, right OT, left frontal, right frontal) at the oddball frequency (i.e., 2 Hz). We conducted a 2 (category: within, cross) × 2 (hemisphere: left, right) repeated-measure ANOVA in the OT and frontal ROIs separately. It revealed significant main effects of color category (*F* (1, 16) = 5.52, *p* = 0.032, 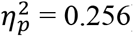) and hemisphere (*F* (1, 16) = 6.35, *p* = 0.023, 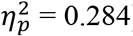) in OT ROIs (Fig. 3A). The interaction effect was not significant (*F* (1, 16) = 0.21, *p* = 0.653, 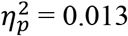). In frontal ROIs, the effect of color category reaches marginal significance (*F* (1, 16) = 4.42, *p* = 0.05, 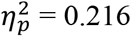). Neither the effect of hemisphere (*F* (1, 16) = 0.70, *p* = 0.417, 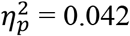) nor the interaction effect was significant (*F* (1, 16) = 2.358, *p* = 0.144, 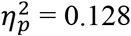). These results suggested that the cross-category condition elicited significantly greater responses than the within-category condition at the oddball frequency over bilateral occipitotemporal regions. This category effect was weaker in the frontal regions.

**Fig.3.**
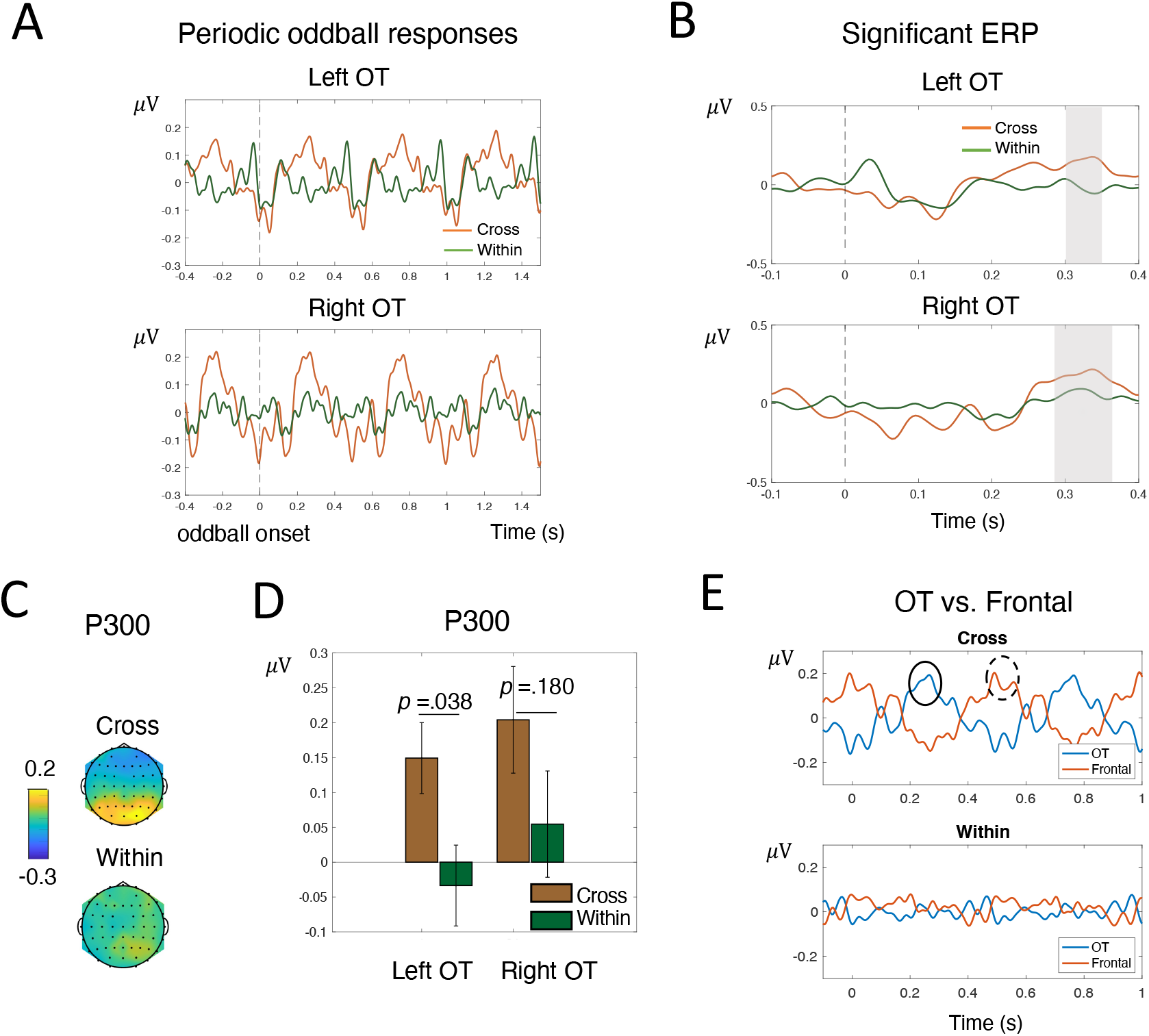
Results of time domain analyses. (A) EEG signals evoked by oddball stimuli over four oddball cycles. (B) Average waveforms after oddball onset in the left OT (top panel) and right OT (bottom panel) in two conditions (red: cross-category; blue: within category). Gray areas indicate time windows significantly greater than zero in the cross-category condition (cluster-based permutation test against zero, *p* < 0.05). (C) Topographic map showing average responses of the P300 (300 ms - 350 ms) in two conditions. (D) Average amplitudes of P300 (300 ms – 350 ms) in the left and right OT regions in two conditions. Error bars represent 95% CI. (E) Periodic EEG signals evoked by oddballs in the OT and frontal regions. In the cross-category condition, the oddball stimuli evoked significant responses in the OT region around 270 ms (solid circle) and in the frontal regions around 500 ms (dashed circle).

One of the major concerns in color category studies is that whether the observed category effect is a genuine “categorical” effect or due to low-level color differences (for review, see Siuda-Krzywicka et al., 2019a). To tackle this issue, we assessed the spatial distribution of the category-selective responses (cross-category minus within-category) at the oddball frequency and compared it with the EEG responses to within-category color stimuli (Fig. 3B). The EEG responses to within-category oddballs are supposed to reflect low-level factors. According to Fig. 3B, we can see that the low-level color responses are peaked in more posterior sites, while the category-selective responses are peaked in more anterior areas. Thus, we believe that the observed category effects in the frequency domain result from cross-boundary changes rather than low-level perceptual factors.

### 3.3 EEG: time domain

Fig.3A presents the response waveform over bilateral OT regions in two conditions, time-locked to the onset of the oddball stimuli, over four oddball cycles. We assessed how the responses to the within- and cross-category differences at the oddball frequencies unfold over time. One-sample cluster-based permutation t-tests against zero revealed significant time windows for both left and right OT regions in the cross-category condition (Fig.3B, left OT: ~ 300 ms – 350 ms; right OT: ~ 290 ms – 370 ms). In contrast, the within-category condition did not show any significant time window. There were no significant differences between these two conditions in the two frontal ROIs.

To take a deeper look at the category effect in the OT areas, we then computed the mean amplitudes of P300 (300 ms – 350 ms) for the within- and cross-category conditions and compared their differences (Fig.3C & Fig.3D). The cross-category condition exhibited significantly greater response in P300 than the within-category condition in the left ROI region (*t* (16) = 2.27, *p* = 0.038, Cohen’s *d* = 0.549). The cross-category condition also exhibited greater amplitudes in P300 in the right ROI region, but the difference didn’t reach statistical significance (*t* (16) = 1.40, *p* = 0.180, Cohen’s *d* = 0.340). The results suggested that EEG signals associated with categorical color coding can be observed in the visual areas approximately 300 ms after stimulus presentation. To examine whether the effect originates from processes within the visual areas or from top-down modulation of high-level cognition (e.g., attention), we extracted the average activity over the OT region (pooling P4, P6, P8, PO4, PO8, P3, P5, P7, PO3, PO7) and the average activity in the frontal region (pooling F1, F2, F3, F4, F7, F8, AF3, AF4, AF7, AF8). As shown in Fig. 3E, the frontal region exhibited large activities in the within-category condition, about 500 ms after the onset of oddball stimuli (marked by the dotted circle). This activity was later than the P300 response in the OT region (marked by the solid circle). Therefore, it is likely that the P300 response in the OT region indeed reflect processes within the ventral visual system rather than from top-down modulation to the visual cortex from high-level cognitive (e.g., attention) processing.

## 4. Discussion

The current study adopted a frequency-tagging-based EEG paradigm to probe whether our brain automatically categorizes briefly presented colors. We showed that the EEG response to cross-category oddballs at the frequency where the oddball stimuli were presented was significantly larger than the response to within-category oddballs. This finding suggested that the categories of color stimuli were likely to be automatically encoded, as the color stimuli were only briefly flashed and irrelevant to the participants’ task. The ERP results showed that cross-category oddballs elicit significant responses at around 300 ms, but this effect was not observed for within-category oddballs.

We found that the brain can encode color categories even when color stimuli were presented rapidly (100 ms SOA) and when top-down feedback processing from frontoparietal areas is minimized by backward masking. It suggested that the categorical color representation is likely to originate from the visual system. This result is consistent with the notion that category effects occur during the perceptual stage. There are two main streams of studies supporting that category effects occur at the perceptual stage. First, category effects were observed in pre-linguistic children (Franklin, Drivonikou, Bevis, et al., 2008; Ozturk, Shayan, Liszkowski, & Majid, 2013; Yang et al., 2016). Second, some EEG studies have found that category effects are associated with early ERP components (Athanasopoulos et al., 2010; Clifford et al., 2010; Fonteneau & Davidoff, 2007; Holmes et al., 2009; Maier & Rahman, 2018; Mo et al., 2011; Thierry et al., 2009; Zhong et al., 2015). It should be noted that the present study found a late ERP component (around 300 ms) associated with color categories. By comparing the response latencies in the occipitotemporal and frontal regions, we found that at around 300 ms, the encoding of color categories in the occipitotemporal region may still reflect local processes within the visual hierarchy. The feedback process from the high-level cortex has not been involved yet, and it takes about 500 ms for neural activity to extend from the occipitotemporal region to the frontal region. Previous studies using the RSVP paradigm found that when visual stimuli are presented at high speed, neural activity in higher-level visual areas that process categorical information is delayed (Mohsenzadeh et al., 2018). This may be the reason for the relatively late appearance of category-selective EEG signals in the occipitotemporal region in our study. Previous EEG studies often make inference of the stage of color categorization based on the latency of ERPs. We believe that this inference has a premise that the visual processing of the brain is not interrupted by the following stimuli. Unlike previous studies, our study strictly limited the processing time of each color stimulus in the brain, intervening in the normal visual process. Therefore, the observation of significant differences in within- and cross-category conditions on the P300 component should not be interpreted solely based on ERP latency as evidence that the processing of color categories occurs in the post-perceptual phase. In addition, the current results do not rule out the possibility that top-down modulation of color perception could get involved in a later stage. The top-down view suggests that category effects may be due to the top-down modulation of low-level color perception by high-level cognitive processes (e.g., language, attention) and that the fine-grained and continuous color perception space is transformed into categorical encoding. While the current study found that the visual areas can represent color categories without top-down influence, this may not be the only way the brain encodes color categories. When stimulus presentation time is not limited, the processing of color categories may involve the high-level frontoparietal areas (e.g., Brouwer & Heeger, 2013; Siok et al, 2009; Sun et al., 2021).

Centered on the universalism-realism debate, the lateralization of category effects has been one of the critical topics in previous studies. The results of frequency domain analyses revealed that the effect of color categories was comparable between the left and right ROIs, showing no evidence of lateralization effects. However, the results from the time domain were a little ambiguous. The cross-category condition exhibited significantly greater responses in P300 than the within-category condition in the left ROI but not in the right ROI, which seems to suggest lateralized category effects. However, the patterns of the category effects in the two ROI regions were actually similar (Fig.4C). We then compared the size of the category effects (cross-category minus within-category) between these two ROIs with paired t-tests, which showed no significant differences. Overall, we did not find robust evidence of lateralized category effects (Holmes & Wolff, 2012; Siuda-Krzywicka et al., 2019b; Sun et al., 2021).

Finally, our study demonstrates that the frequency-tagging-based EEG paradigm is a sensitive tool for exploring categorical color encoding in the brain, as we observed significant categorical effects for briefly presented color stimuli. In previous studies, only Maier and Rahman (2018) used a brief SOA (94 ms) similar to the current study. They found that the color category effects occurred in the early perceptual stage and was related to the P1 and N2 components. However, different from our study, Maier and Rahman (2018) adopted the attentional blink paradigm, and the subjects need to respond to two targets (T1 and T2) in the experiment. This task requirement may influence the processing of the target and the non-target. Studies have shown that attention and prediction modulate ERP components related to visual processing such as P1 (Baumgartner et al., 2018; Mohr et al., 2020). Thus, the P1 component found in Maier and Rahman (2018) may be affected by attention. The current study does not require the subjects to respond to color stimuli, thus ruling out the possibility that the observed effect may be reflect attentional requirement of an explicit task.

One limitation of our study is that the color samples are pre-determined and identical to all participants. There are small individual differences in color naming (Forder et al., 2017), and thus color samples based on the group level might obscure the category effects. Future studies should take individual differences into consideration and select color samples based on each individual’s category boundaries.

## 5. Conclusion

This study adopted the frequency-tagging-based EEG paradigm and found that the brain categorizes color stimuli presented with a SOA of 100 ms and backward masked. We suggested that the brain can encode color categories automatically when top-down feedbacks from frontoparietal areas are blocked. Therefore, categorical color perception is likely to originate from the visual cortex.

## Credit author statement

**Mengdan Sun**: Conceptualization, Methodology, Formal analysis, Writing – original draft, Visualization; **Xiaoqing Gao:** Conceptualization, Methodology, Writing – review & editing, Fund Acquisition.

